# Beyond Fixation: detailed characterization of neural selectivity in free-viewing primates

**DOI:** 10.1101/2021.11.06.467566

**Authors:** Jacob L. Yates, Shanna H. Coop, Gabriel H. Sarch, Ruei-Jr Wu, Daniel A. Butts, Michele Rucci, Jude F. Mitchell

## Abstract

Virtually all vision studies use a fixation point to stabilize gaze, rendering stimuli on video screens fixed to retinal coordinates. This approach requires trained subjects, is limited by the accuracy of fixational eye movements, and ignores the role of eye movements in shaping visual input. To overcome these limitations, we developed a suite of hardware and software tools to study vision during natural behavior in untrained subjects. We show this approach recovers receptive fields and tuning properties of visual neurons from multiple cortical areas of marmoset monkeys. Combined with high-precision eye-tracking, it achieves sufficient resolution to recover the receptive fields of foveal V1 neurons. These findings demonstrate the power of free viewing to characterize neural response while simultaneously studying the dynamics of natural behavior.

**Highlights:**

- We introduce a free-viewing paradigm for studying neural mechanisms of visual processing during active vision
- Receptive fields (RFs) and neural selectivity in primary visual cortex (V1) and area MT can be extracted during free-viewing in minimally-trained subjects
- Novel high-resolution eye tracking in this context supports detailed measurements of receptive fields in foveal V1

## Introduction

The investigation of perception and cognition in systems neuroscience has relied extensively on frameworks that employ highly controlled, repeatable animal behavior. Although it has long been recognized that to understand the function of neural systems, we must also understand their operation in the context of the natural behaviors for which they evolved, experimental approaches have favored simpler, less natural behavioral paradigms to maintain rigor. This has been no truer than in visual neuroscience, where parametrically controlled stimuli and parametrically controlled behavior have been the gold standard.

All animals with image-forming eyes acquire visual information through eye movements (1), which shape the visual input by constantly changing the retinal image and its temporal dynamics (2). However, standard characterizations of neural processing of vision, to date, require stabilization of the subject’s gaze – either through anesthesia/paralytics (3,4) or trained fixation on a central point (5) (Figure 1a) – or they simply ignore eye movements entirely (6). Even experiments that involve active components of vision – such as covert attention, or the planning of saccadic eye movements – primarily involve analyses during instructed saccades and fixation on a point (7–9).

**Figure 1.**
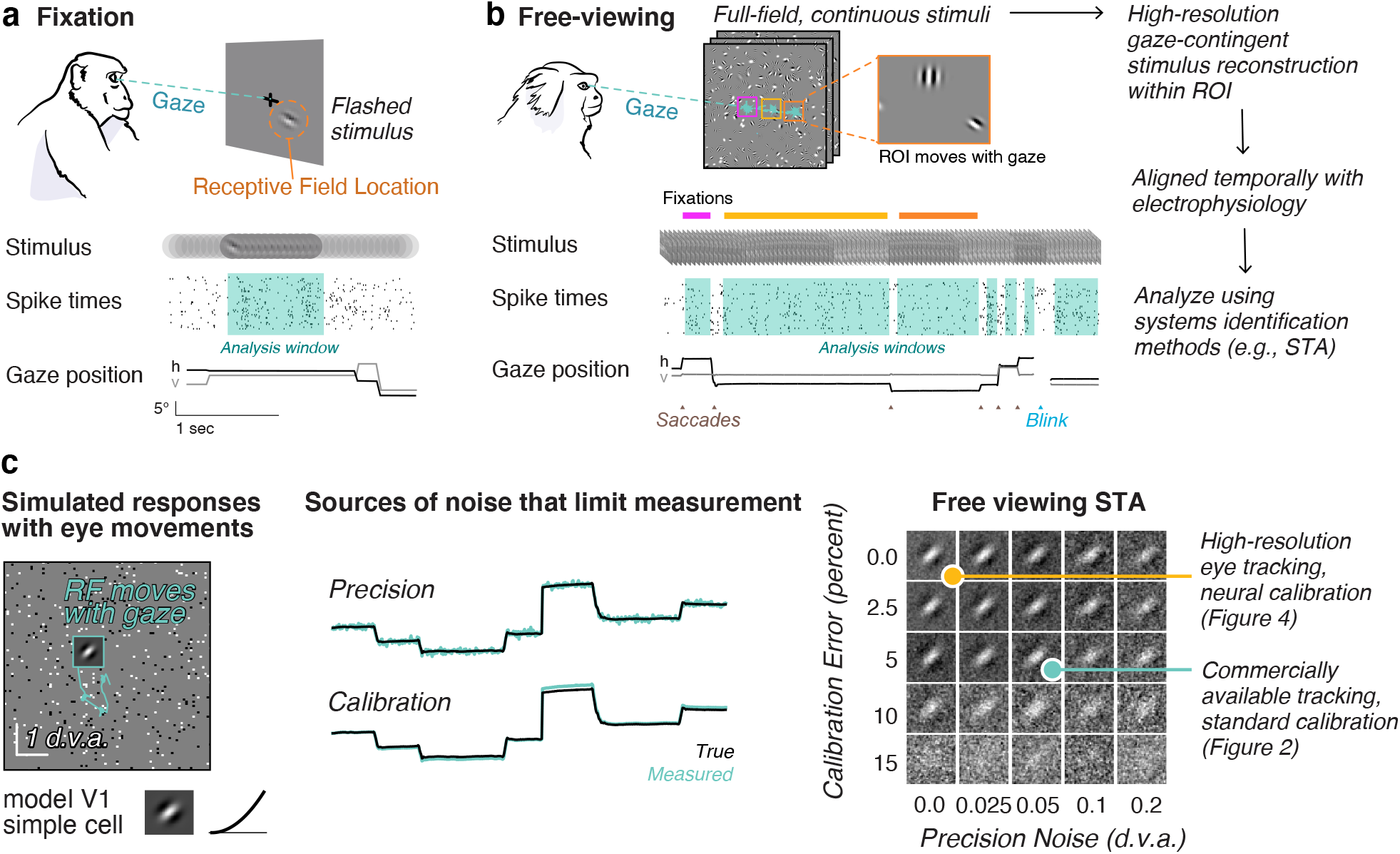
Free viewing paradigm and gaze-contingent neurophysiology. **a**. Conventional fixation paradigm with flashed stimuli. Spike times are aligned to stimulus onset and analyzed during a window during fixation. **b**. Free viewing: subjects freely view continuous full-field stimuli. Shown here is dynamic Gabor noise. All analyses are done offline on a gaze-contingent reconstruction of the stimulus with a region of interest (ROI). Analysis windows are extracted offline during the fixations the animal naturally produces. **c**. Simulation demonstrates how uncertainty in the gaze position (due to accuracy and precision) would limit the ability to map a receptive field. (Left) A model parafoveal simple cell (linear receptive field, half-squaring nonlinearity, and Poisson noise) moves with the gaze. Example gaze trace shown in cyan. RF inset is 1 d.v.a. wide. (Middle) errors in precision are introduced by adding Gaussian noise to the gaze position. Errors in calibration are introduced by a gain factor from the center of the screen. (Right) The recovered RF using spike-triggered averaging (STA) on a gaze-contingent ROI. Of course, with zero precision noise or calibration error, the STA recovers the true RF. Adding either source of noise degrades the ability to recover the RF, however, some features are still recoverable for a wide range of noise parameters at this scale. RFs that are smaller or tuned to higher spatial frequencies require high resolution eye tracking. The yellow and cyan dots indicate two levels of accuracy that are explored in figures 2 and 5, respectively.

While these conventional paradigms have given us a standard and highly successful model of early visual processing (10,11), it is unknown how well those results generalize to describe natural visual conditions. Relatively few labs have attempted to study visual processing during natural eye movements (12,13), and none have been able to interpret neural responses with respect to detailed visual processing in the presence of natural eye movements. Recent experimental work has demonstrated that eye movements modulate neural selectivity substantially in many brain areas (14,15). Moreover, eye movements are how visual input is normally acquired. Recent work suggests this process is fundamental in formatting the visual input to facilitate normal vision (16–18). Furthermore, fixation paradigms come with a substantial cost in our understanding of visual processing: the visual stimulus the subject is looking at (the fixation point) is not the stimulus under study (19,20)

To study natural vision without any loss of detail or rigor, we have developed a suite of integrated software and hardware tools to characterize neural selectivity during natural visual behavior, and do so at a resolution that exceeds standard fixation paradigms. Our approach, “free viewing”, lets subjects look wherever they please within the visual display. We perform all analyses on a gaze-contingent reconstruction of the retinal input. Although previous studies have “corrected” for small changes in eye position by shifting the stimulus with the measured or inferred center of gaze, this has only been attempted for small displacements of the stimulus during instructed fixation(21–23). Our approach differs in that the subjects are free to explore the visual scene, and therefore, there are no training requirements.

Previous attempts at free viewing have faced three main obstacles: 1) the computational requirements to recover receptive fields from full-field stimuli, 2) limitations in eye tracker precision, and 3) errors resulting from eye-tracker calibration. Here, we show that combining image-computable neural models with correction for eye position from commercially available eye tracking is sufficient to recover receptive field size and tuning in free-viewing animals. Additionally, we introduce a novel high-resolution eye tracker for non-human primates and offline calibration using measured neurophysiology to give sufficient resolution to study visual processing in foveal neurons of primary visual cortex.

We demonstrate this approach here in marmoset monkeys. Marmosets are small new-world primates with homologous visual architecture to larger primates (24) and similar eye-movement statistics (25). They are increasingly used as a model for neuroscience because of their similarity to humans and benefits for genetic tools (26). However, marmosets are limited in their ability to fixate for prolonged periods. Our approach circumvents this issue, making both standard and new neural characterization approaches possible in marmoset, and also resulting in a higher data-throughput per animal: generating more data per unit time than fixation paradigms. It also provides an opportunity for rigorous study of visual neuroscience in species where fixation paradigms may be impractical (such as ferrets, tree shrews, and rodents).

We apply the free-viewing approach to recover receptive field properties to both primary visual cortex (V1) where neurons can have extremely high spatial and temporal resolution, and area MT, a higher visual area specialized for motion processing. Additionally, with novel high-resolution eye tracking it is possible to recover finescale spatial receptive field structure of neurons in the central degree of vision (the foveola) for the first time.

## Results

### The free-viewing paradigm

To study natural vision in untrained animals, we depart from conventional approaches that stabilize the subject’s gaze behaviorally with a fixation cross. Instead, we present full-field natural and artificial stimuli in 20 second trials while monitoring eye position and neural activity. Figure 1b illustrates the free viewing approach: because the retina moves with the eyes and visual neurons have receptive fields in a retinal coordinate frame, we must correct for changes in eye position to correctly represent the visual inputs to neurons. The relevant stimuli for a set of neurons can be recovered offline using a gaze-contingent region of interest (ROI) that moves with the eyes. Once the stimulus is reconstructed within the gaze-contingent ROI, conventional analysis tools can be used. In the following sections, we describe the successful application of this approach to recordings from V1 and MT of 3 marmoset monkeys (Callithrix Jacchus; 2 males, 1 female).

### Retinotopy and selectivity in V1 during free-viewing paradigms

A considerable barrier to using free-viewing paradigms prior to this work is limitations in eye-tracking. Figure 1c demonstrates the effect of eye tracking limitations by simulating the responses of a model V1 simple cell with a receptive field that moves with the eyes and adding common sources of noise. Of course, if the experimenter perfectly recovered the true eye position, they could recover the receptive field because the gaze-contingent input would be identical the input with stabilized gaze. However, real eye trackers have noise that effects the precision of their measurements (top trace, precision). Eye trackers must also be calibrated, which is only as accurate as the subject’s ability to fixate on points on the screen presented during the calibration procedure, which has inherent error associated with it (bottom trace, calibration). Adding these sources of noise affects the ability of the experimenter to recover an RF from the free viewing approach (Figure 1c, right panel).

A second obstacle to free-viewing is computational limitations in processing full-field high-resolution stimuli. A standard monitor today has 1920 × 1080 pixels. Generating artificial stimuli at high frame rates and full resolution is now possible with gaming graphics processing units (GPUs) and procedurally generated stimuli can be reconstructed offline at full resolution for part of the screen.

In this section, we show that full-field sparse noise stimuli and commercially available eye tracking (Eyelink 1000) can be used to recover the size, location and tuning of receptive fields (RFs) in V1 during free viewing. The sparse noise stimulus allows us to efficiently estimate RF locations over a large portion of the visual field, which is often all that is required for further targeting neurons with behavioral paradigms, but also can be used to further target analyses with high-resolution stimuli within an ROI.

We present sparse noise consisting of flashed dots or squares in random positions on each frame (Figure 2a) during free viewing and use a gaze-contingent analysis to align the stimulus to retinal coordinates. We move a grid with the location of gaze on each frame (Figure 2a). As the grid moves with the eyes, we then average the stimulus luminance within each grid location on each video frame. This can be computed rapidly using the position and sign of the dots that are present on each frame of our noise stimulus. Our initial ROI is 28 × 16 degrees of visual angle (d.v.a.) with 1 d.v.a square bins, centered on the gaze location. This window covers a large portion of central vision, including all possible retinotopic locations in our recording chamber (and an equal area in the opposite hemifield).

**Figure 2.**
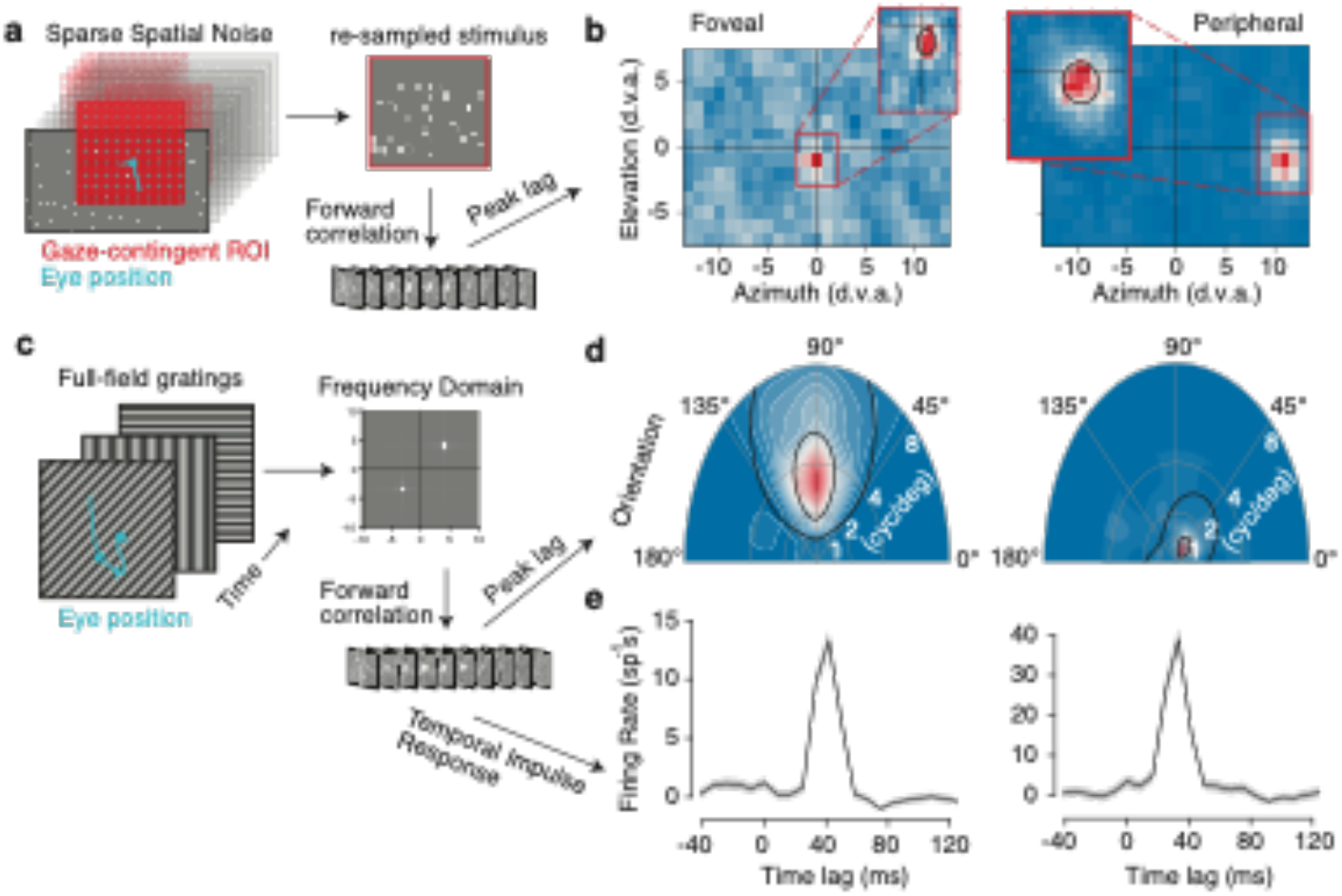
Receptive field mapping and feature tuning in V1. **a.** Retinotopic mapping approach. A gaze-contingent grid with 1° spacing is used to estimate the RF. **b.** Spatial RF for foveal and peripheral example units. Insets indicate analysis at a .3° grid size. The black line is a Gaussian fit at 2 standard deviations. **c.** Subspace reverse correlation procedure for mapping tuning using full-field gratings. **d.** Example joint orientation-frequency tuning maps for the units in panel b. Black lines indicate 50 and 75% contour lines from a parametric fit (methods). **e.** Temporal impulse response in spikes per second for preferred gratings measured with forward correlation.

We used regularized linear regression (methods) to estimate the spatiotemporal RF. As illustrated by the spatial response profile at the peak temporal lag (Figure 2b), we can identify receptive fields at a coarse spatial scale (1 d.v.a. bins) for two example neurons: one in the fovea and one in the periphery.

We then re-define a new ROI centered on the RF and run the same binning and regression procedure at a finer spatial scale within the new ROI, using .3 d.v.a. bins (Figure 2b, insets). Spatial RFs were typically recovered with less than 5 minutes of recording time. The median recording time to recover spatial retinotopic maps was only 2.86 minutes and the range was 1.33 to 21.85 minutes (n=44 sessions). We then fit a 2D gaussian to the finescale spatial map to recover the RF location and size. Though it is possible to train marmosets to perform conventional fixation tasks (25,27), much of that time would be unusable for analysis due to breaks between the trials and limited trial counts. Using the free-viewing approach here results in a substantial gain in the total analyzable neurophysiology data over fixation paradigms (Supplemental Figure 1).

The free-viewing analysis was sufficient to recover spatially selective RFs in 410 of 739 recorded units from marmoset V1, and demonstrated a highly comparable relationship between eccentricity and size of RFs as reported from previous literature with anesthetized marmosets (Supplemental Figure 2).

We also measured visual feature selectivity during free viewing using sinewave gratings. We presented full-field gratings that were updated randomly on each frame (Figure 2c) and performed subspace reverse correlation (28) yielding the spatial-frequency RF for the same two example units, plotted in polar coordinates where angle represents stimulus orientation and radial distance represents spatial frequency (Figure 2d). This analysis produces selective responses in 437 of 739 recorded units and worked well across the visual field. The subspace reverse correlation also gave temporal response functions consistent with known V1 temporal response profiles (Figure 2e). The median recording time used for grating receptive fields was 10.84 (ranging from 7.08 to 43.33) minutes (n = 53 sessions). The resulting distribution of preferred orientations was possible across a range of eccentricities (Supplemental Figure 2) and was comparable to previous reports from macaque V1 (27). Thus, the feature tuning of neurons in V1 can be measured during free viewing with short recording times in minimally-trained marmosets using commercially-available eye tracking and standard calibration.

### Free-viewing approach recovers receptive field properties in area MT

The validity of this approach is not limited to simple visual features that drive primary visual cortex, but can generalize to other features and higher level visual areas. We demonstrate this here by measuring motion-selective RFs for neurons recorded from area MT during free-viewing. Extra-striate area MT is a higher-order visual area with the vast majority of neurons exhibiting exquisite tuning to retinal motion (29). To measure motion-selective RFs, we adapted the sparse noise stimulus described above to include motion. noise stimulus described above to include motion. Rather than simply appearing and disappearing on each frame, noise dots drifted for 50 ms in one of 16 directions. Using the same gaze-contingent analysis window, we converted the spatiotemporal stimulus into separate horizontal and vertical velocity components (Figure 3a). This spatiotemporal velocity stimulus was then used as the input to a Linear Nonlinear Poisson (LNP) model of the MT neuron spike trains (methods).

**Figure 3.**
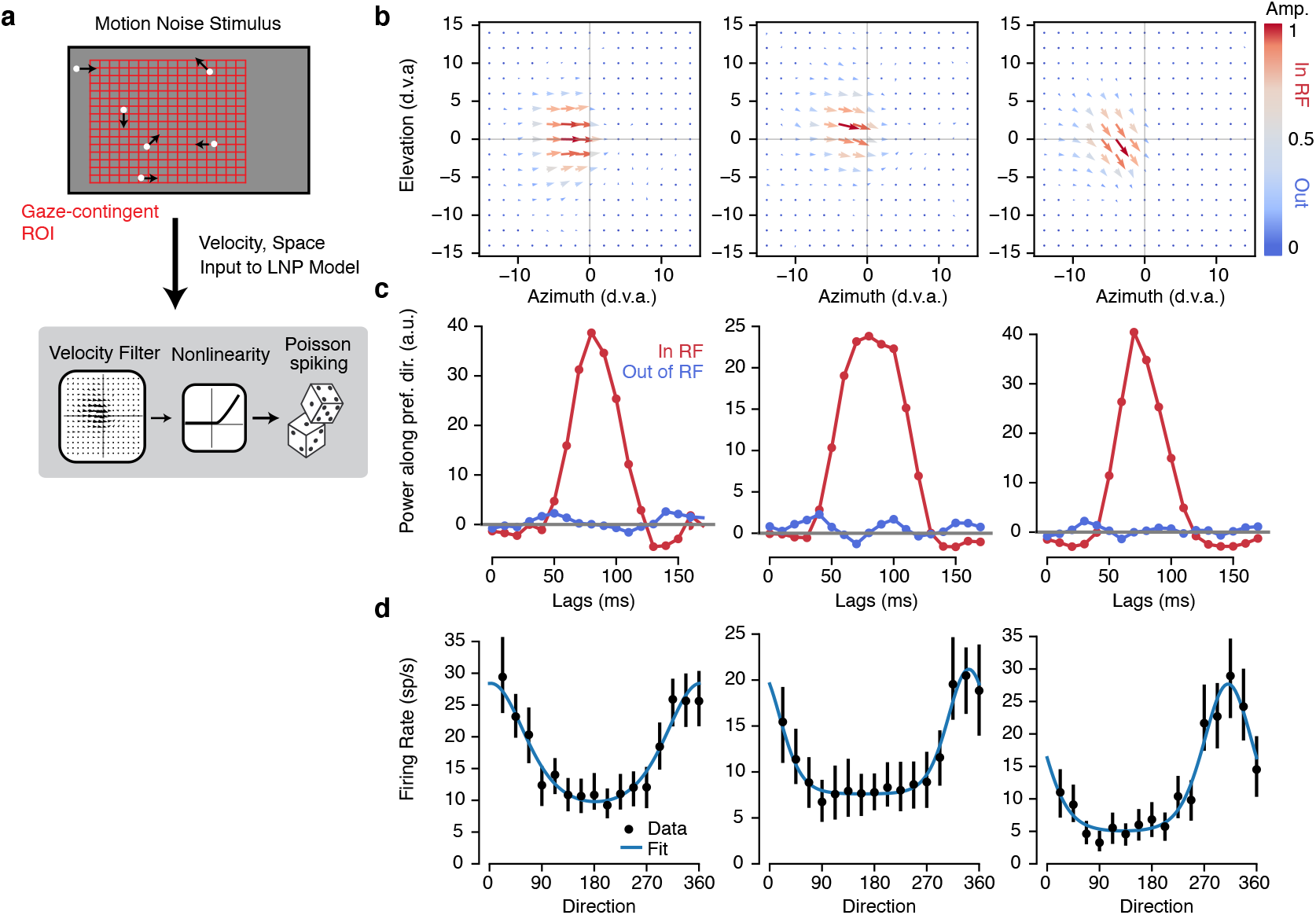
Receptive field mapping and tuning in MT. **a.** Sparse motion-noise stimulus was converted into a spatiotemporal velocity stimulus with separate horizontal and vertical velocities using a gaze-contingent grid with 2° spacing. This downsampled stimulus is used to estimate the receptive field (RF) using linear nonlinear Poisson model (LNP). **b.** The spatial map at the peak lag of the spatiotemporal velocity RF from the LNP fits is shown as a vector plot for three example units. Color indicates the vector amplitude (ranging from 0 to 1, with gray at 0.5). **c.** The temporal impulse response was measured both in and out of the RF by projecting the (unnormalized) vector at the maximum and minimum amplitude of the spatial RF on the preferred direction (unit vector). The three plots correspond to the three example units in b. **d.** Tuning curves were measured by masking the stimulus with half of the max of the spatial RF and computing the firing rate at the peak lag for each direction shown. Error bars are 95% confidence intervals measured with bootstrapping and blue lines are fits with a von mises function.

### Free-viewing approach recovers receptive field properties in area MT

The validity of this approach is not limited to simple visual features that drive primary visual cortex, but can generalize to other features and higher level visual areas. We demonstrate this here by measuring motion-selective RFs for neurons recorded from area MT during free-viewing. Extra-striate area MT is a higher-order visual area with the vast majority of neurons exhibiting exquisite tuning to retinal motion (29). To measure motion-selective RFs, we adapted the sparse noise stimulus described above to include motion. Rather than simply appearing and disappearing on each frame, noise dots drifted for 50 ms in one of 16 directions. Using the same gaze-contingent analysis window, we converted the spatiotemporal stimulus into separate horizontal and vertical velocity components (Figure 3a). This spatiotemporal velocity stimulus was then used as the input to a Linear Nonlinear Poisson (LNP) model of the MT neuron spike trains (methods).

The LNP model trained on gaze-contingent velocity stimuli recovered spatiotemporal velocity RFs for MT neurons (Figure 3b). We found detailed spatiotemporal measurements of the velocity selectivity of MT neurons, which we decomposed into spatial maps of direction selectivity (Figure 3b), temporal selectivity (Figure 3c), and overall motion tuning (Figure 3d, see methods). The MT neurons in our sample were well fit by von Mises tuning curves (mean rsquared = 0.90 +- 0.013, n=21 units). This highlights that full-field stimuli can be engineered to target complex feature selectivity and that regression-based analyses can recover detailed spatiotemporal measurements of that selectivity during unconstrained visual behavior.

### High-resolution eye tracking for detailed 2D spatiotemporal receptive fields in the fovea

No previous studies have accurately recovered the full spatiotemporal receptive field structure of V1 neurons in primate foveal regions. This gap is not due to negligence, but rather reflects limitations in the accuracy of eye tracking. Even anesthetized, paralyzed monkeys exhibit drift in eye position over the course of an experiment. Further, the conventional approach to obtain accurate RF estimates using fixation paradigms obscures study of foveal vision because the center of gaze is occupied by the fixation point as the stimulus. Another limitation to all previous studies is that fixation is imperfect, with continuous eye drift and fixational eye movements, which are substantial and would limit precision if uncorrected (Supplemental Figure 3). The free viewing approach provides an opportunity to directly stimulate foveal vision to recover high resolution RFs if it used in conjunction with sufficiently accurate eye-tracking. Here, we apply free viewing with high resolution (both in terms of accuracy and precision) eye tracking to measure detailed receptive fields in the fovea of free-viewing marmosets.

To obtain precise measurements of gaze position, we adapted a recently developed video eye tracker (30) for use with marmosets. The digital Dual Purkinje Imaging (dDPI) eye tracker uses a digital CCD camera, IR illumination and GPU processing to track the 1^st^ and 4^th^ Purkinje images achieving a 0.005 degree precision (RMS of noise measured with an artificial eye) and is precise enough to measure and correct for fixational drift and microsaccades (Supplemental Figure 3).

To achieve full-resolution receptive fields in the fovea, we center an ROI on the retinotopic location of the recorded neurons (as in Figure 2) and then reconstruct the full stimulus (pixel by pixel) for every frame that within that ROI (as in Figure 1b). Beyond flashed spatial dots or gratings, we also presented trials in which the free viewing background consisting of flashed Gabor and Gaussian stimuli of varying phase, orientation, and spatial scales. For 5 foveal recording sessions, we reconstructed every frame of the experiment within a 70 × 70 pixel gaze-contingent ROI. This was done for every stimulus condition so that we had a gaze-contingent movie of the stimulus at the projector refresh rate (240 Hz).

While the dDPI tracker used in the current study provide high precision position signals, they must be properly calibrated to align them with visual space to obtain an accurate estimate of the actual eye position. To calibrate high-resolution eye-trackers, previous studies in humans use a two-stage calibration procedure, where the human subjects adjust their own calibration parameters in a closed loop (31). As our marmosets were unlikely to perform self-calibration without extensive training, we developed an offline calibration method using V1 physiology directly. Briefly, we fit a convolutional neural network (CNN) model of V1 that included a recalibration of the eye tracker to optimize the model fits to gaze-contingent neural activity across the recorded session. Specifically, the CNN predicts the spiking response of the entire population of simultaneously recorded units given the spatiotemporal gaze-contingent stimulus movie, which is reconstructed based on the eye tracker calibration identified by the CNN (Figure 4a). One critical aspect of this method is that we obtain better fits to the eye tracker calibration when we use recordings from larger V1 populations. Therefore, using recordings with high-density arrays as used in this current study represents a distinct advantage.

**Figure 4.**
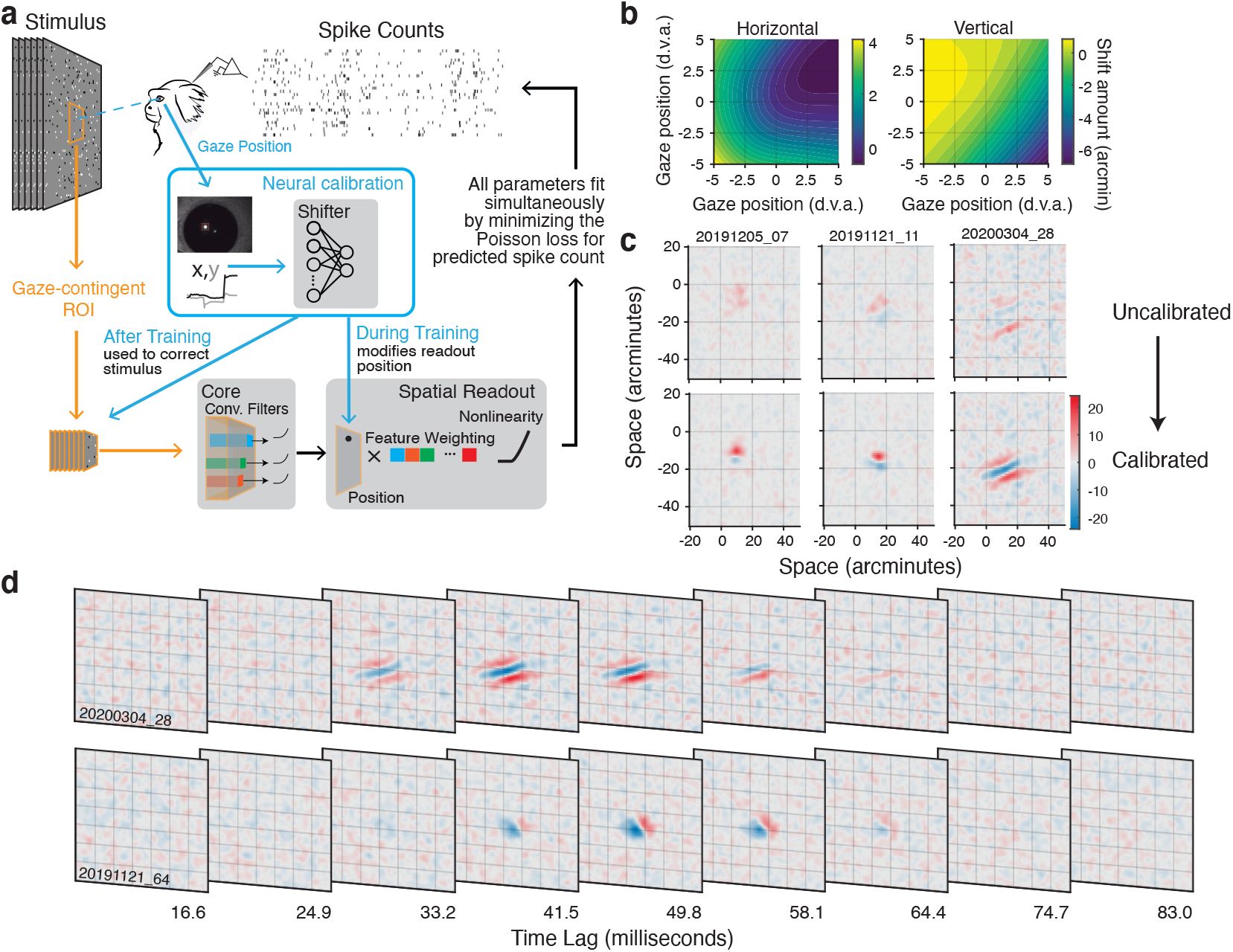
Neural eye-tracker calibration and high-resolution foveal receptive fields. **a.** Convolutional Neural Network (CNN) architecture used to calibrate the eye tracker. The gaze-contingent stimulus within the ROI is processed by the nonlinear subunits of the convolutional “Core”. The “Spatial Readout” maps from the core to the spike rate of each neuron with a spatial position in the convolution and a weighted combination of the feature outputs at that position. This is passed through a static nonlinearity to predict the firing rate. The “Shifter” network takes in the gaze position on each frame and outputs a shared shift to all spatial readout positions during training. All parameters are fit simultaneously by minimizing the Poisson loss. After training, the shifter output is used to shift the stimulus itself so further analyses can be done on the corrected stimulus. **b.** Calibration correction grids for horizontal and vertical gaze position are created using the output of the shifter network. These are used to correct the stimulus for further analysis. **c.** Spatial receptive fields before (top) and after (bottom) calibration for three example foveal units demonstrates the importance of calibration for measuring foveal RFs. **d.** Example foveal spatiotemporal RFs. Grid lines are spaced every 10 arcminutes. The axes bounds are the same as in c.

The key component of the CNN is a 2-layer “shifter” network which corrects the calibration of our eye tracker. The shifter network takes the gaze position on each frame as input and produces a *shared* shift to the spatial position of *all* units being reconstructed in our recording session (Figure 4a). Any errors in the behavioral calibration of the eye tracker will manifest as shifts in the RF locations for all units as a function of where the marmoset is looking, which will be learned by the shifter network. Such corrections to the output of the dDPI eye tracker end up being smooth functions of spatial position, as illustrated by the output of the shifter network for an example session visualized as a function of gaze position (Figure 4b). As was typical of our sessions, the shifter network produced small shifts (maximum 7 arcmin, median shift 1 arcmin) across a 10 d.v.a. range of gaze positions. These calibration matrices (CMs) dictate how to correct the initial experimental calibration. Across repeated fitting, the shifter network produced highly reliable CMs, suggesting that the shifter was measuring systematic changes in RF position as a function of the monkey’s gaze position, likely both a result of small errors in the initial calibration established by the experimenter, and systematic deviations from linearity of the calibration.

Once an accurate calibration was established, we can reconstruct the detailed retinal input to the receptive field at foveal precision. We compared the linear RF computed with spike-triggered averaging (STA) with and without the improved CM correction. Figure 4c shows example foveal RFs measured with and without correction. The foveal RFs recovered from these sessions were 22.56 [21.63,23.79] arcminutes away from the center of gaze and were 10.76 [10.25,11.14] arcminutes wide. The examples in Figure 4c illustrate that the shifter network is necessary to make detailed measurements in the fovea, as reflected by the RFs measured with and without correction.

RFs post correction had amplitudes that were significantly larger than without correction (median ratio = 1.65 [1.55, 1.75], p < 7.9 × 10^-16, Wilcoxon signed rank test, zval=8.054, signed rank=3741).

Our recordings from marmoset V1 are the first detailed 2D spatiotemporal measurements of foveal cortical processing. Figure 4D shows example single neuron spatiotemporal RFs. Across neurons, we found a range of spatial and temporal receptive field properties that are consistent with several classic findings of simple and complex cell receptive field structure in V1, provided a substantial miniaturization for the spatial scale. These preliminary findings decisively demonstrate the power of the free viewing methodology when combined with high-resolution eye tracking and neural eye tracker calibration. They also open a new avenue of research for examining foveal scale visual representations not only in V1 but also other visual area along the ventral processing stream which specialize in higher acuity object vision.

## Discussion

We introduced a free-viewing paradigm for visual neuroscience. This approach is higher-yield (per unit recording time) than fixation-based approaches and yields measurements of spatial RFs and feature tuning in minimally-trained animals. It works with standard commercially available eye trackers for standard descriptions in V1 (Figure 2) and MT (Figure 3). We also demonstrated this paradigm can be extended to study foveal V1 neurons by introducing a high-resolution video eye tracker based on the dual Purkinje method and a calibration routine based on the output of V1 neurons. Combined with free viewing, these methods support state-of-the-art in the measurement of 2D spatiotemporal receptive fields of neurons in the foveal representation of V1 (Figure 4). Thus, the three major advantages to free viewing, especially when combined with high-resolution eye tracking, are: 1) application to untrained animals, 2) increased usable data per unit recording time, and 3) measurement of foveal visual processing. Further, although we only analyzed epochs of stable fixation between saccades in the present study, the free-viewing paradigm will also have major advantages study of the role of eye movements during visual processing.

Free viewing is amenable to any animal model with almost no training requirements. Here, we applied this approach to marmosets. In the last decade, mice have emerged as a popular model for visual neuroscience (32). Despite the fact that mice move their eyes in a directed manner (33,34), the spatial position of gaze is rarely accounted for in neurophysiological studies of visual cortex in mouse. A notable exception to this, Lurz et al., 2020, used a similar CNN model to the one we employ to model mouse V1 (35). Although they did not use the learned shifter network to calibrate the gaze and measure spatial RFs as we did here, the model improvement from shifting the stimulus implies they would see similar improvements in RF measurements in mouse V1 after correcting the stimulus as we did. Similarly, the free viewing paradigm is a potentially promising future direction to expand rigorous visual neuroscience to animal models with higher acuity and smaller RFs than mice, but without the ability to perform trained fixation (such as ferrets and tree shrews). This type of paradigm will also support a direct comparison of visual processing and modulatory signals in multiple species, such as the role of locomotion in visual processing for non-human primates.

Offline gaze-contingent analysis of neural data during free-viewing opens the possibility of studying neural computations in a range of natural visual behaviors and exceeds the resolution set by fixation studies. Although, previous studies have corrected for small changes in eye position by shifting the stimulus with the measured or inferred center of gaze, this has only been attempted for small displacements of the stimulus during instructed fixation (21–23). Our approach differs in that the subjects are free to explore the visual scene, and therefore, both the calibration and the displacements must be accounted for. Importantly, our use of visual cortex to calibrate the eye tracking differs from previous approaches such as neural-based eye tracking (22) in that all of the temporal dynamics of gaze are directly measured by a physical eye tracker that is independent of receptive field properties, as opposed to being dynamically inferred from neural activity. The CNN used neural activity to improve the calibration of the eye tracker, but after that remained fixed in subsequent analyses of receptive fields. The added precision also makes it possible to examine the role of fixational drift and eye movements, an essential component of vision (16). Further studies could assess, for example, whether RFs in V1 are explicitly retinotopic or dynamically shift to account for small fixational movements as proposed by recent theoretical work (36). And as illustrated in Figure 4, this approach affords the opportunity to examine foveal receptive fields in primate V1 for the first time. Despite its paramount importance for human vision, almost nothing is known about neural processing in the foveal representation.

Finally, one limitation of our approach is by letting the eye’s move freely, there are no longer repeats of the same stimulus condition, which is one of the main workhorses of systems neuroscience. However, fixational eye movements preclude that reality, even where it has been used previously (23). With the development of higher speed cameras and video displays, it will soon be possible to stabilize retinal images during free viewing, thus affording more precise control of the stimuli input to visual neurons than previously possible in fixation paradigms. Examining natural behaviors that lack fixed repetitions is possible with comparable rigor as conventional approaches as shown here, when using appropriate neural models to fit the responses to natural stimuli. In near future, the application of neural models during natural behavior will finally allow us to gain deeper insight into the dynamics of neural processing in natural contexts.

## Acknowledgements

National Institutes of Health grants K99EY032179 (JY), R01EY18363 (MR), and 1R01EY030998 (JM). JY was an Open Philanthropy fellow of the Life Sciences Research Foundation. National Science Foundation grants GRF DGE1745016 & DGE2140739 (GS). We thank Amelia Wattenberger for help with figure 1.

## Methods

### Surgical procedures

Data were collected from 3 adult common marmosets (Calithrix jacchus; one female and two males). All surgical and experimental procedures were approved by the Institutional Animal Care and Use Committee at the University of Rochester in accordance with the US National Institutes of Health guidelines. At least one month prior to electrophysiological recordings, marmosets underwent an initial surgery to implant a titanium headpost to stabilize their head during behavioral sessions. Surgical procedures for the headpost procedures were identical to those described previously (27).

A second surgery was performed under aseptic conditions to implant a recording chamber. For the chamber implantation, marmosets were anesthetized with intramuscular injection of Ketamine (5-15 mg/kg) and Dexmedetomidine (0.02-0.1 mg/kg). A 3D-printed chamber (http://www.protolabs.com) was then attached to the skull with metabond (http://www.parkell.com) over coordinates guided by cranial landmarks. A 3×4 mm craniotomy was then drilled within the chamber (http://www.osadausa.com). The dura was slit and exposed tissue was covered with a thin layer (< 2mm) of a silicone elastomer (World precision instrument, https://www.piinc.com) as in Spitler et al., 2008.

### Electrophysiological Recordings

Electrophysiological recordings were performed using multisite silicon electrode arrays. The arrays consisted of 1-2 shanks, each containing 32 channels separated by 35 or 50 μm. The electrode arrays were purchased from NeuroN-exus (http://www.neuronexus.com) and Atlas Neuro Engineering (https://www.at-lasneuro.com). We recorded from neurons using a semi-chronic Microdrive system. We adapted the EDDS Microdrive System (https://microprobes.com) for use with silicone arrays and to be removable. Our chamber and drive designs are available online (https://mar-molab.bcs.rochester.edu/resources.html). A reference wire was implanted under the skull at the edge of the chamber. The electrode arrays were lowered through the silicone elastomer and into brain using a thumbscrew.

Data were amplified and digitized at 30kHz with Intan headstages (Intan) using the open-ephys GUI (https://github.com/open-ephs/plugin-GUI). The wideband signal was highpass filtered by the headstage at 0.1 Hz. We corrected for the phase shifts from this filtering (Okun, 2017). The resulting traces were preprocessed by common-average referencing and highpass filtered at 300Hz. The resulting traces were spike sorted using Kilosort or Kilosort2. Outputs from the spike sorting algorithms were manually labeled using ‘phy’ GUI (https://github.com/kwikteam/phy). Units with tiny or physiologically implausible waveforms were excluded.

### Eye-tracking and saccade detection

Gaze position was monitored using one of two eye-trackers. 28 sessions did not involve high-resolution measurements in the foveal representations. Eye position was sampled at 1000Hz using an Eyelink 1000 (SR Research).

For 30 high-resolution sessions, a custom digital Dual-Purkinje Imaging system (dDPI) was used. The dDPI uses a collimated IR beam (ThorLabs) a dichroic mirror (Edmunds) and samples 0.4 Megapixel images of the eye at 539 frames per second (DMK 33UX287; The Imaging Source). Custom CUDA code running on a gaming GPU (GTX 1080Ti; Nvidia) performs the algorithm described in (30) to extract the eye position. Briefly, the 1^st^ and 4^th^ Purkinje images (P1 and P4) were identified and tracked using a two-stage process. The first stage is to find the region of interest (ROI) for each. The camera image is downsampled by a factor of 4 and P1 is found by thresholding the 8-bit image at 200 and calculating the center of mass of the pixels exceeding the threshold. The ROI for P4 was found via template matching on the down-sampled frame. Following the initial ROI finding stage, the center of each Purkinje image was calculated using the full-resolution image within each ROI by center of mass for P1 and radial symmetric center for P4 (cite).

Methods for calibrating both eye-trackers before a behavioral session were identical to those described previously (25,27). Briefly, this procedure sets the offset and gain (horizontal and vertical) of the eye-tracker output manually. The calibration was refined offline using a bilinear regression between the eye position during a detected fixation and the nearest grid target (within a 1 degree radius) during the calibration routine.

Saccadic eye movements were identified automatically using a combination of velocity and acceleration thresholds as described in (37). The raw eye position signals were resampled at 1 kHz, and horizontal and vertical eye velocity signals were calculated using a differentiating filter. Horizontal and vertical eye acceleration signals were calculated by differentiation of the velocity signals using the same differentiating filter. Negative going zero crossings in the eye acceleration signal were identified and marked as candidate saccades. These points correspond to local maxima in the eye velocity signal. Eye velocity and acceleration signals were then examined within a 150 ms window around each candidate saccade. Candidate saccades were retained provided that eye velocity exceeded 8° /s and eye acceleration exceeded 2000°/*s*^2^. Saccade start and end points were determined as the point preceding and following the peak in the eye velocity signal at which eye velocity crossed the 10°/s threshold.

### Visual Stimuli and Behavioral Training

For all V1 recording sessions, visual stimuli were presented on a Propixx Projector (Vpixx) with a linear gamma. All stimuli were generated in Matlab (the Mathworks) using the Psychtoolbox 3 (38). Stimulus and physiology clocks were aligned and synchronized using a Datapixx (Vpixx) following the method described in (PLDAPS paper). Stimulus code is available online at (https://github.com/jcbyts/MarmoV5).

### Foraging task

All visual protocols besides the static natural images were run simultaneously with a “foraging” paradigm where marmosets obtained a small juice reward (marshmallow water) for fixating small (0.5 – 1.0 d.v.a diameter) targets that would appear randomly in the scene. Reward was granted any time the marmosets kept their gaze within a specified radius of the center position of the target for more than 100ms. Targets consisted of either oriented Gabor patches (2 cycle/deg) or marmoset faces that were taken from photos of the colony. Marmosets will naturally look at faces (25) and these were used to encourage participation in the forage paradigm. The position of the targets was generated randomly near the center of the screen (either drawn from a 2D Gaussian at the center or from an annulus with a 3 d.v.a. radius) to encourage the gaze to stay near the center of the screen where eye-tracking accuracy and precision are highest. The amount of reward was titrated based on the subject’s performance to ensure they did not get too much marshmallow in a single session (5-10μl per reward).

### V1 retinotopic mapping and receptive field size

Retinotopic mapping stimuli consisted of full-field randomly flashed high-contrast circles or squares (referred to as “dots” from here on). Each dot was either white or black, and appeared at a random position anywhere on the screen. Across sessions, the dot-size and number of dots per frame varied, but were fixed within a session.

Offline gaze-contingent retinotopic mapping was performed in a two-stage process using regularized linear regression (39). First, we estimated the RF at a coarse resolution and then re-sampled the stimulus at a finer resolution within an ROI centered on the result of the first stage. The coarse resolution RF was created by re-sampling the dot stimulus on a gaze-contingent grid. This rectangular grid, *G*_*x,y*_, consisted of 405 locations spaced by 1 d.v.a. from −14 to 14 d.v.a along the azimuthal axis and −8 to 8 d.v.a. of elevation. The resampled gaze-contingent stimulus is a vector *X*(*t*)at frame *t* and was calculated by summing over the dots on each frame

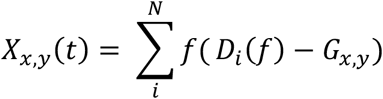

where *D*_*i*_(*f*) is a vector of the position of an individual dot on frame *t* and *f* is a function that returns a vector of zeros with a 1 at the grid location where the dot was centered. This method is fast and does not require regenerating the full stimulus at the pixel resolution. Additionally, by summing the number of dots within each grid location, this analysis ignored the sign of the dot (“black” or “white”) relative to the gray background, which was designed to target cells that exhibited some phase invariance (i.e., complex cells). We found that ignoring sign generated more robust retinotopic mapping results with fewer datapoints.

We estimated the spatiotemporal receptive, *Ksp*, of each unit, *i*, by using regularized linear regression between the time-embedded gaze-contingent stimulus *X* and the mean-subtracted firing rate, *R*, of the units binned at the frame resolution.

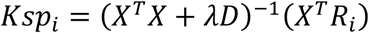

Where *D* is a graph Laplacian matrix corresponding to spatial and temporal points in *X* and *λ* is a scalar that specifies the amount of regularization. *λ* was chosen using cross-validation. This measures the spatiotemporal receptive field (RF) in units of spikes per second per dot. We then repeated this processes at a 0.3 d.v.a grid size centered on the RF location recovered from the coarse stage. We found the RF location by thresholding *Ksp* at 50% of its max an used the matlab function regionprops to find the centroid and bounding box. We scaled the bounding box by 2 and re-ran the regression analysis to estimate the final RF. We then fit a 2D Gaussian to the spatial slice at the peak lag using least-squares with a global search over parameters and multiple starts. We calculated the Euclidean distance between the RF centroid location and the mean of the Gaussian fit and normalized that by the eccentricity of the mean (distance from 0,0). We converted the fitted covariance matrix to RF area using the following equation: Units were excluded if the mean shifted by more than 0.25, meaning that the fitting procedure produced a Gaussian that was not centered on the RF centroid. This resulted in 410/739 units with measurable spatial RFs.

### V1 Tuning

To measure the neurons’ selectivity to orientation and spatial frequency, we flashed full-field sinewave gratings. Gratings were presented at 25% contrast and were either drawn from the Hartley basis (Ringach 2002) or were parameterized using a polar grid of orientations and spatial frequencies. On each frame, only one grating was presented and up to 50% of the frames were a blank gray background. We represented the frequency space on a polar basis. The basis consisted of 8 evenly spaced von mises functions for orientation, and 4 nonlinearly stretched raised cosine functions for spatial frequency. This basis served to convert stimuli collected with the Hartley set and the polar grid to the same space. We measured the grating receptive field *Kgr* for each unit using regularized linear regression (as described above for the spatial mapping).

To measure the tuning curve of the units, we fit a parametric model of the form

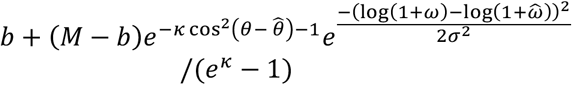

Where θ is the orientation, *ω* is spatial frequency, 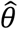 and 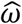 are the orientation and spatial frequency preference, respectively; *b* is the baseline firing rate, *M* is the maximum firing rate, *κ* and σ scale the width of the orientation and spatial frequency tuning. This parametric form combines a normalized von Mises tuning curve for orientation that wraps every π and a loggaussian curve for spatial frequency tuning (each normalized to have a maximum of 1 and a minimum possible value of 0). We converted the dispersion parameters into bandwidths as the full-width at half height of each curve. Tuning curves were fit using nonlinear least squares (lsqcurvefit in matlab).

### MT velocity receptive fields and direction tuning

MT mapping stimuli consisted of sparse dot motion noise. Every video frame contained up to 32 white dots that were 0.5 degrees in diameter. Each dot was either replotted randomly or moved at 15 degrees/s in one of 16 uniformly-spaced directions with a life-time of 5 frames (50 ms with frame rate at 100 hz). Marmosets performed the foraging task while this motion-noise stimulus ran in the background.

To calculate the RFs, the dot displacement on each frame transition was split into horizontal and vertical velocity components at each spatial location on a gaze-contingent grid with 2 d.v.a. wide bin size. This produced two gaze-contingent spatiotemporal stimulus sequences of the same form as described for V1 retinotopic mapping methods separate for horizontal and vertical velocities. Velocity receptive fields were measured by fitting a Poisson Generalized Linear Model (GLM) to the spike trains of individual MT units. The parameters of the GLM include the RF of the unit and a bias parameter to capture baseline firing rate. The RF parameters were penalized with to support spatial smoothness and sparseness using the same Graph Laplacian penalty used for retinotopic mapping and an L1 penalty. Example fitting and analysis code is available at https://github.com/jcbyts/neureye/.

To measure the temporal integration of MT RFs, we first computed the “preferred direction vector” of the unit as the weighted average of the recovered RF at the peak lag. We then found the spatial location with the largest amplitude vector and calculated the projection of the RF direction vector at that location onto the preferred direction vector for all time lags. We repeated this for the spatial location with the smallest amplitude vector.

To measure the direction tuning curves, we masked the stimulus spatially at every location greater than half of the max of the spatial RF and counted the number of dots drifting in each direction on each frame. We then calculated the direction-triggered firing rate of each unit through forward correlation between the directions on each frame and the firing rate, normalized by the number of dots shown. The tuning curve was taken to be the value for each direction at the peak lag. Error bars were computed using bootstrapping and correspond to 95% confidence intervals. We fit a von Mises function to the firing rate *R*.

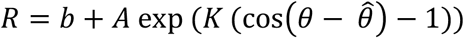

Where *b* is the baseline firing rate, *A* is the amplitude, *K* is the bandwidth and 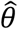 is the preferred direction.

### Full resolution stimulus reconstruction

Stimuli were reconstructed by playing back the full experiment. All randomized stimuli were reconstructed using stored random seeds and the replayed frames were cropped within the gaze-contingent ROI using using Psychtoolbox function Screen(’GetImage’). Randomized stimuli included flashed dots described earlier, as well as flashed Gabor noises that will be described in detail below. All high-resolution analyses operated on this reconstruction. For the 5 sessions we analyzed high-resolution RFs, the width and height, *w* × *h*, of the ROI was 70 × 70 pixels, where each pixel subtends 1.6 arcminutes.

### Neural eye tracker calibration

The network used for calibrating the eye consisted of three parts: a *core* neural network that forms a nonlinear basis computed on the stimulus movies, a *readout* for each neuron that maps from the nonlinear features to spike rate, and a *shifter network* that predicts RF shifts using the measured gaze position. The architecture here was based on the networks that have previously been successful for modeling V1 responses (35,40). The neural network machinery in this application enabled us to optimize the weights in the shifter network and establish correction grids that shift the eye tracker’s output into a more accurate estimate of eye position.

The CNN core consists of a single layer of 16 subunits. Each subunit has a 19 × 19 2D convolutional filter. Time was embedded in the stimulus using the channel dimension. The output of the convolutional filters are passed through a nonlinearity modeled after divisive normalization (41). Specifically, the filter outputs are passed through an exponential linear unit (ELU) nonlinearity plus one and then divided by a weighted sum

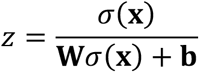

Where σ(**x**) is the 16 × *w* × *h* output of the convolutional ELU units, **W** is a 16 × 16 matrix constrained to be positive that specifies the divisive drive from each unit to each unit, and **b** is a 16 × 1 vector that specifies the semi-saturation constant. **W**σ(**x**) is a weighted sum over the channel dimension of 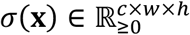.

The readout maps the nonlinear features from the core to the spike rate of each neuron through an instantaneous affine transformation. To reduce the number of parameters and make the readout interpretable with respect to space, we used a factorized readout where each neuron, *i*, has a vector of feature weights, *v*_*i*_ (that correspond to the 16 output channels from the core), and a spatial readout that specifies a position in the convolutional output the combines across the core output using bilinear interpolation (35,40). Following the approach in (35), the *x*, *y* position were sampled from a Gaussian with mean μ_*i*_ and covariance Σ_*i*_ for each neuron. This reduces the spatial readout from 70 × 70 per neuron to 5 parameters per neuron.

The shifter network consists of a 2-layer network with 20 hidden units with SoftPlus activation functions in the first layer and 2 linear units in the second. It takes the measured gaze position on each frame that was used for the stimulus reconstruction and outputs a horizontal and vertical shift that affects *μ*_*i*_ for all neurons. The output of the shifter network was constrained to be 0 for the gaze position 0,0. All parameters were learned simultaneously using stochastic gradient descent with the Adam optimizer with weight decay (42,43). All stimulus sets were used for training the parameters of the model. A Pytorch implementation with examples is available at (https://github.com/jcbyts/neureye).

The shifter calibration matrices were constructed by passing in a grid of potential eye positions from −5 to 5 d.v.a centered on the center of the screen. These 2D correction grids were then used to correct the stimulus on each frame for gaze positions within that region. Specifically, the measured eye position on each frame corresponds to a location in the correction grid. That location corresponds to an amount of shift. We used bilinear interpolation to map from gaze position to shift amount in the grid. The use of correction grids maps the two-layer shifter network into an interpretable format and these can be used across multiple stimulus sets to correct both the measured eye position and the gaze contingent stimulus reconstruction.

### High-resolution receptive fields

Receptive fields were recovered for the highresolution stimuli using spike-triggered average (STA) on the pixels of reconstructed stimulus. The stimuli used for these RFs was either a sparse noise or Gabor noise stimulus. The sparse noise consisted of .1 d.v.a diameter black or white dots positioned randomly on each frame with a density of 1.5 dots/deg2 on each frame. The Gabor noise consisted of multiscale Gabor patches with carrier frequencies ranging from 1 to 8.5 cycles per degree and widths (standard deviation of Gaussian) ranging from 0.05 to 0.132 d.v.a with a density of 2 Gabors/deg2. The STA was computed 12 lags at 120 frames per second

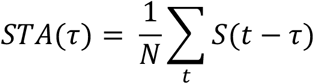

Where *N* is the number of spikes, *t* is the frame index for each spike, *S* is the stimulus frame, and τ is the lag. STAs were z-scored for visualization with the same normalizing constants before and after calibration.

---

**Supplemental Figure 1.**
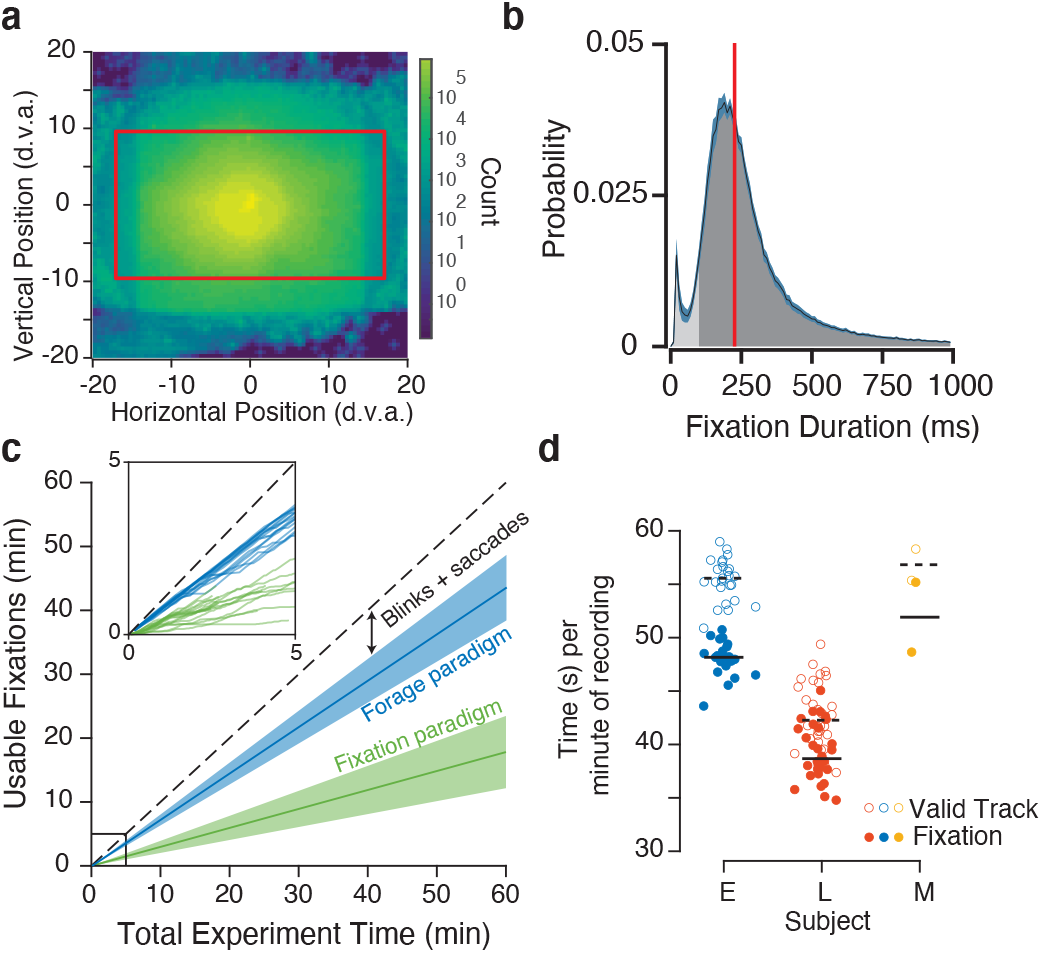
Eye movements during free viewing and usable fixation time. **a.** Distribution of gaze position for three marmosets during foraging as a heat map. Most time is spent within the central 10 degrees. The circular density at the center is reflective of the annulus from which the forage targets were generated. The red box indicates the size of the projector screen for high-resolution eye tracking sessions. **b.** Distribution of fixation durations during free viewing for 3 marmosets. The dark shaded region indicates the fixations that were included for analyses. Red vertical line indicates the median fixation duration. **c.** Analysis of the usable fixation time clipping out fixations from free viewing compared to using a fixation paradigm in marmosets. The inset shows measured times for N fixation and N free viewing sessions in the same marmosets. **d.** Fixation time per minute of recording during free viewing for three marmosets (E, L, M). Open symbols indicate the time with valid eye-tracking (eyes open, track good) and filled symbols are the time spent fixating.

**Supplemental Figure 2.**
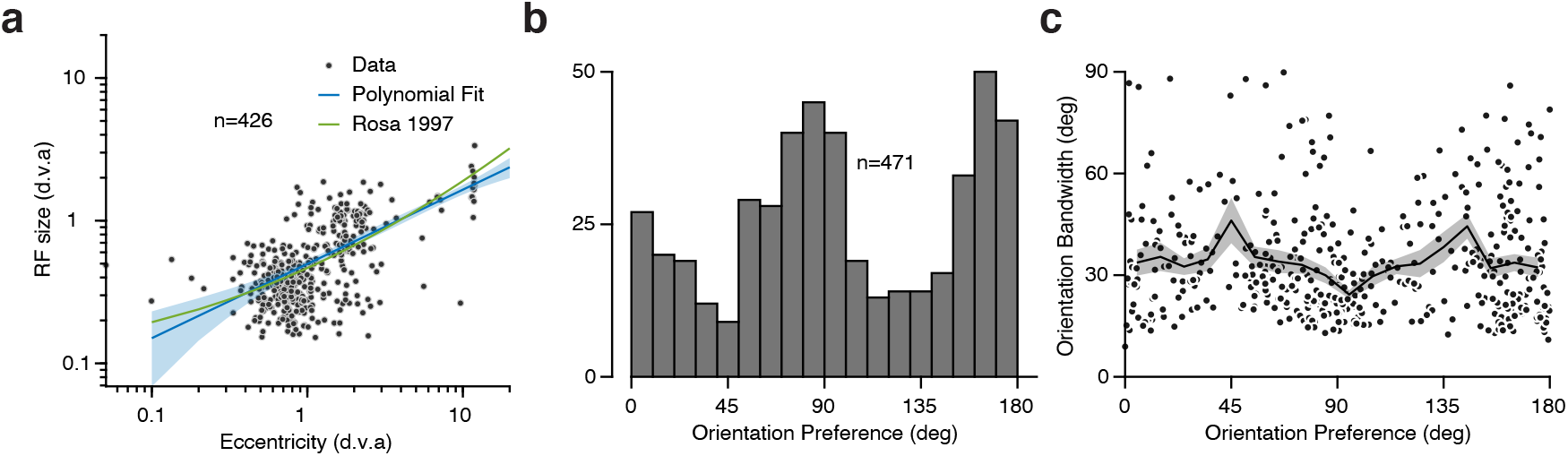
Receptive field size and tuning in V1. **a.** RF size (square-root of area) as a function of eccentricity (n = 426). Blue line is 2^nd^-order polynomial fit to the log eccentricity. Shaded area depicts 95% confidence intervals. Green line is the same fit using reported parameters from Rosa et al., 1997. **b.** Distribution of preferred orientations measured using parametric fit to the spatial frequency reverse correlation. **c.** Tuning bandwidth (full-width at half max) as a function of orientation preference. Black line is a running average in 10° bins. The shaded region corresponds to 95% confidence intervals measured with bootstrapping.

**Supplemental Figure 3.**
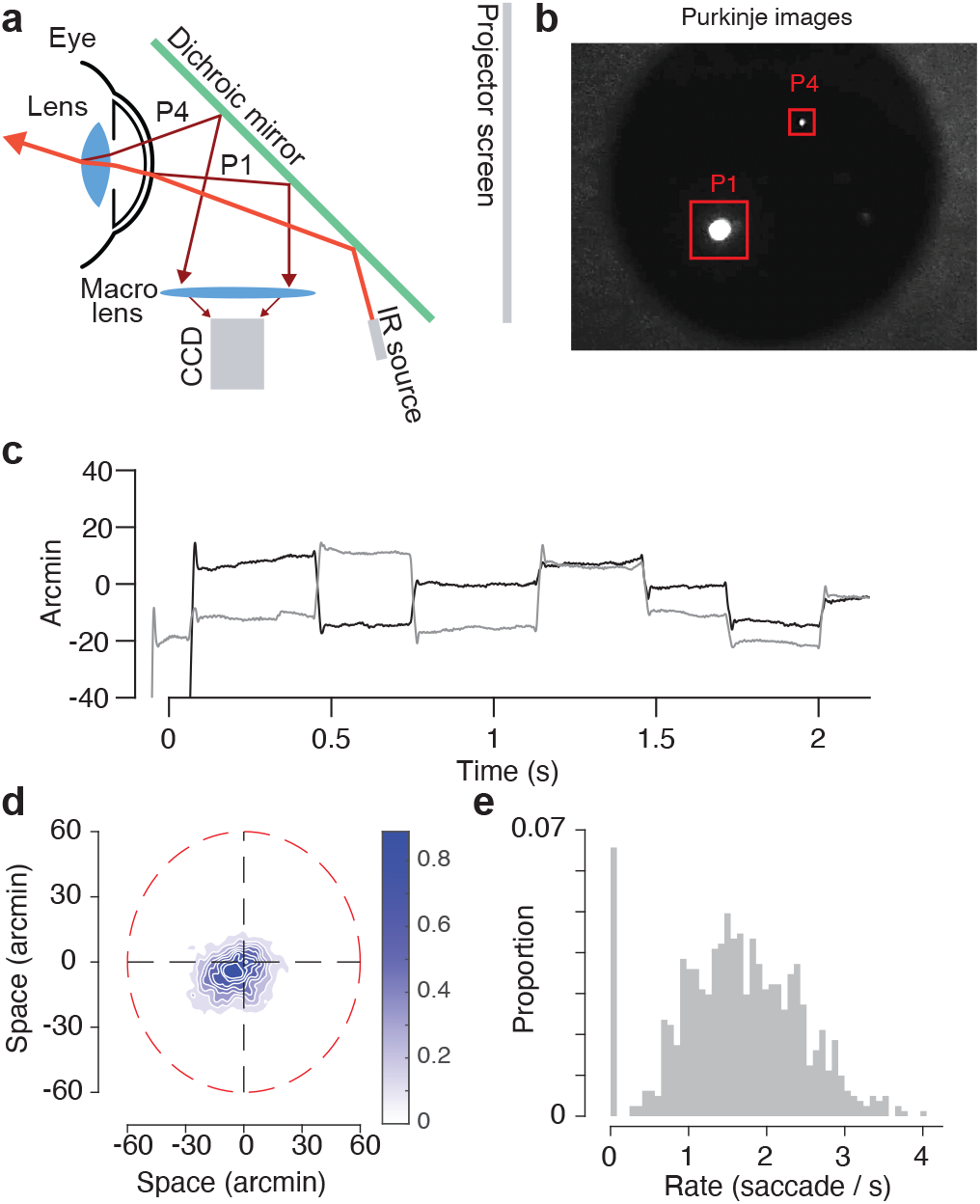
High resolution eye tracking. **a.** Schematic of the digital Dual Purkinje Imaging (DDPI) system. **b.** Example frame of the pupil with the first and fourth Purkinje images identified. All tracking was performed online using a GPU. **c.** Example horizontal (black) and vertical (gray) gaze position measurements during a fixation trial. This trial was picked because it had many microsaccades within the fixation window. **d.** Gaze position scatter during fixation. Red dashed line indicates the fixation window that was used online. **e.** Histogram of microsaccade rate during fixation.

